# Teacher Attitudes towards Research-Based Classroom Exercises and Intensive Virtual Summer Professional Development

**DOI:** 10.1101/2022.07.01.498491

**Authors:** Miroslav Nestorovic, Gregory J. Gage

## Abstract

A study based on a survey questionnaire was conducted with 55 teachers across multiple high schools in the United States to understand their perceptions about their attitudes and confidence towards using research-based classroom exercises, which models they use to develop such exercises and their availability and preferences for an intensive summer professional development course. Our preliminary results indicate that while teachers are comfortable teaching research-based classroom activities, they are none-the-less very interested in paid professional development courses on teaching research practices to students (79%), more than two-thirds (69%) would be willing to devote at least one hour a week during the school year and three-quarters (75%) would be willing to spend 2-3 hours a week or more during the summer.

## Introduction

A recent (Nov, 2020) report on the Next Generation Science Standards (NGSS, 2013) adoption from the NGSS Early Implementers Initiative stated that in the 7 years after the standards were released, teachers still faced significant challenges adopting phenomenon-based lesson plans, and that one should “not underestimate how significant the shift is for teachers” to implement three-dimensional learning (Tyler et al., 2020). The transition to new pedagogy requires sustained professional learning, and the recommendation is for teachers to participate in programs that teach NGSS by doing the NGSS. These experiences can shift the teaching approaches and instructional materials to support coherence from the student’s perspective (Reiser, et al. 2017).

Nowhere is the lack of NGSS-aligned K12 education more obvious than in neuroscience. Understanding the brain remains a profound and fascinating challenge that captivates the scientific community and the public alike, and the field of neuroscience is highly integrated across many fields of science (with cross-cutting concepts). The lack of effective treatments for most neurological disorders makes training the next generation of neuroscientists, physicians and engineers a key concern. Despite this, K12 students receive little education in neuroscience (Myslinski, 2022; Hopkins, 2021; Gage, 2019; Dekker & Jolles, 2015; Fulop & Tanner, 2012; Sperduti et al., 2012). This is not only due to the brain’s complexity, but also because the tools and curricula needed to study it are either too expensive or too difficult to use. Given the limited space reserved for neuroscience in NGSS and other state standards, we conducted a qualtrics survey of over 500 teachers to understand if neuroscience could be taught in the classroom (de Freitas et al., 2021). Of the 91 respondents, we found that 4 in 5 teachers (80%) stated they could add neuroscience activities. Of these, 54.5% felt that neuroscience was well supported by their state standards, while 43.2% said they could justify it even if not in state standards. Others mentioned having set times for extra topics (e.g., after standards exams). Over the past decade, our team has produced educational resources by adopting modern neuroscience research techniques into user-friendly and open-source neuroscience curricula suitable for an undergraduate neuroscience lab (Pollak et al., 2019; Nguyen et al., 2017; Shannon et al., 2014; Dagda et al., 2013; Marzullo & Gage, 2012). While some high schools are using these lessons in their program, there has not been a wide adoption of the curriculum in K12. Anecdotally, we found that high school classrooms that use the lessons for students to generate and test their own hypotheses tended to be taught by former research neuroscientists with Ph.Ds who are comfortable discussing the brain and running open-ended experiments. We hypothesize that the issue is that our lessons were developed by Ph.D. neuroscientists and not K12 teachers. What is needed now is to have K12 teachers themselves develop neuroscience lessons so that they feel confident that they understand the material and can see themselves successfully performing the labs in their classrooms.

To address this problem, it is necessary to establish a dialogue across multiple stakeholders in order to understand their perceptions and needs. To begin this process, we conducted a survey of high school teachers across multiple states in the United States, in order to understand teachers’ perceptions and confidence about equipping and empowering them to develop lesson plans that not only deliver neuroscience content, but also overcome practical obstacles of adopting NGSS practices. The aim is to develop best practices for teachers through workshops, lectures, and by virtually simulating a realistic NGSS high school classroom, where several neuroscience hands-on projects are performed by independent groups of students. Our organization has also partnered with the University of Michigan, leading neuroscientists, and industry experts to provide teachers with an academic, educational, and technical context and resources.

## Methodology

We designed a survey to investigate the United States high school teachers’ attitudes towards participating in an in-depth professional development (PD) course, with the goal of building teachers’ self-efficacy in delivering phenomenon-based lessons (Appendix 1). The goal of this survey was to understand teachers’ perceptions about their attitudes and confidence towards the following questions:

- How much experience do teachers have doing scientific research?
- How comfortable do teachers feel about leading phenomenon-based classroom activities where the student acts as a scientist and the outcomes of experiments are uncertain?
- Would teachers be interested in professional development activities to help develop their skills leading independent scientific research in their classroom?
- How many hours a month could teachers be available to participate in paid, virtual (remote) professional development while they are still teaching classes in their home institution and/or when they are done teaching for the summer?
- How interested would teachers be in participating in a virtual paid summer professional development that lasts 10 weeks, a portion of which would start while they were still teaching in their home institution?
- Which methodologies do teachers use to develop Three Dimensional Learning (Scientific Practices, Core Ideas, and Cross-cutting concepts) lesson plans?

All questions were multiple-choice in order to facilitate the quantitative analysis. The demographic information was also collected for statistical purposes. The process used to develop the survey is described in Figure 1.

**Figure 1.**
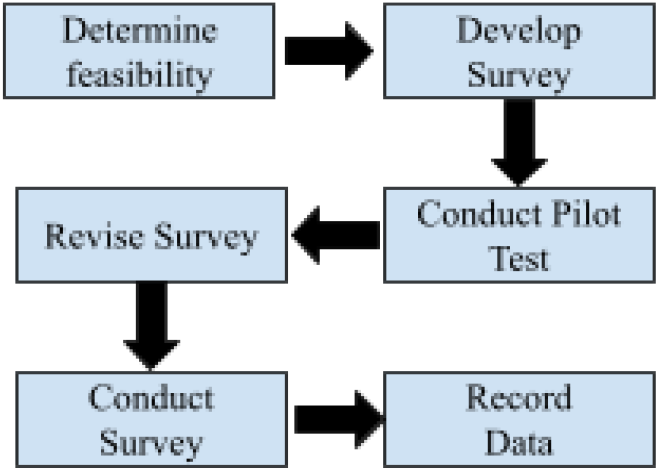
Survey development process

The survey was disseminated using Qualtrics to a pool of 851 teachers who are within the Backyard Brains’ network. Most of these K12 teachers purchased our equipment in the past or provided their email addresses at one of our science or education conference booths.

## Results

We received 55 responses to our survey, most of whom taught at the high school level (82%). Participants taught a variety of STEM subjects including biology (24%), anatomy and physiology (11%), general science (11%), chemistry (9%), engineering (7%), physics (6%) and neuroscience (6%).

Responses to the important question “How much experience do you have doing scientific research?” revealed that 63% of teachers took a research class or worked in a lab in the university or high school. Combined with the results of our previous Qualtrics survey (de Freitas et al., 2021), according to which 67% of participating teachers had been teaching for more than 11 years, this demonstrates that for many teachers it has been a long time since they themselves conducted research.

When asked how comfortable they feel about leading phenomenon-based classroom activities where the student acts as a scientist and the outcomes of experiments are uncertain, 84% of the teachers felt comfortable or very comfortable. Nevertheless, nearly all teachers, 94%, were interested in professional development opportunities to develop their skills leading independent scientific research in the classroom. For this type of professional development 69% of teachers could devote one hour or more a week to participate in paid, virtual (remote) professional development while they are still teaching classes in their home institution. Once teachers are done teaching for the summer, three-quarters (75%) stated that they could devote 2-3 hours a week or more. Out of these, 40% could devote 2-3 hours a week, 13% could devote an hour a day, 4% could devote 2-3 hours a day, and 19% could devote more than 4 hours a day.

Nearly all teachers, 79%, stated that they are interested or very interested in participating in a virtual paid summer professional development course that lasts 10 weeks. Responses also expressed interest in viewing an online weekly lecture series targeted at high school teachers and focused on developing research-based lesson plans (23%) or involving the latest neuroscience research and the brain, done by the University professors (22%).

The most predominant methodologies that teachers use to develop Three Dimensional Learning (Scientific Practices, Core Ideas, and Cross-cutting concepts) lesson plans are the Backward Design model (Identify desired outcomes, determine evidence of learning, plan learning experiences) (19%) and the BSCS 5E instructional model (Engage, Explore, Explain, Elaborate, Evaluate) (16%). 39% of teachers answered that when working on group projects they typically have 5-6 groups in their classroom.

## Discussion

Taken together, these results indicate that there is a significant interest in a 10 week PD course aimed at teaching neuroscience research through the development of phenomena-based lessons. Of the 55 respondents, 79% responded that they would be interested or very interested in attending such a course. Moreover, our survey found that science teachers are willing to implement well-documented phenomenon-based projects in their classrooms.

However, science teachers have very limited experience in performing the research processes they are tasked to train students in NGSS, rarely extending beyond the research-component of their own STEM undergraduate degrees. This can create hesitation to develop phenomenon-based lessons, especially in a field not typically covered in K12.

This can be addressed by recruiting a target teacher population and building their phenomenon-based learning confidence, equipping them with university and pedagogical resources, maker knowledge and mentorship around performing rigorous neuroscience experiments designed for the classroom. Mastering the NGSS curriculum and evaluation can also provide an economic boost for teachers, as problem-based skillsets are widely sought after in districts.

Although this survey is showing promising results, the study has a couple of limitations. First, the participants are familiar with Backyard Brains, most of the teachers had provided their email address at one of the Backyard Brains’ science or education conference booths or purchased our equipment in the past, and may have used our neuroscience classroom products before. Therefore they are subject to biases and other teachers may not be as comfortable teaching research-based classroom activities or as interested in professional development with the goal of leading students’ independent scientific research. Second, the survey sample size is relatively small and the internal consistency was not assessed.

To address these limitations in the future, we expect to increase our sample size beyond the Backyard Brains’ network and to increase the research rigor throughout the survey development process. We also identified opportunities to engage with multiple stakeholders in order to understand their perceptions about the value of neuroscience in K12 and the best ways to conduct research-based classroom exercises in secondary education by addressing the significant challenges that teachers still face in adopting phenomenon-based lesson plans.

Overall, the positive impact of active learning is well documented in STEM education. However, if teachers who teach neuroscience related topics are too disconnected from the underlying principles of phenomenon based learning, they can inadvertently mitigate and remove important elements of neuroscience in the classroom. Thus, we argue that the successful delivery of neuroscience in the classrooms has a direct correlation not only with learning objectives and teaching styles, but that the neuroscience lessons should be developed by K12 teachers themselves, so they feel confident that they understand the material and can see themselves successfully performing the labs in their classroom. Also, a supportive environment is needed that recognizes the value of neuroscience in the schools.

## Acknowledgments

GJG is a co-founder and co-owner of Backyard Brains, Inc., a company that manufactures and sells neuroscience equipment. MN is employed by Backyard Brains, Inc.

## Appendix 1.

The appendix gives all 14 questions and the possible answers from the survey. The answers are presented as blue bars representing the percentages relative to the total number of answers for each question.

Q1. What is the name of the school(s) where you teach? Please, specify: (data not shown)

Q3. What grades do you teach? (select all that apply)

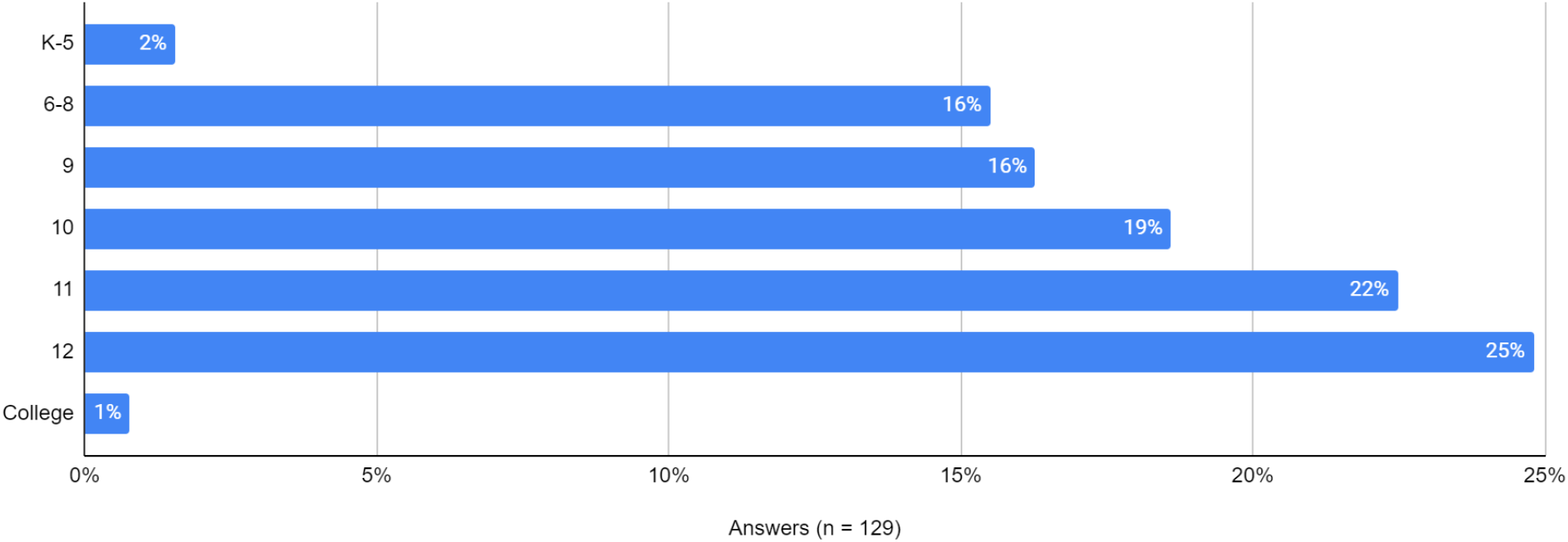

Q4. What courses do you teach? (select all that apply)

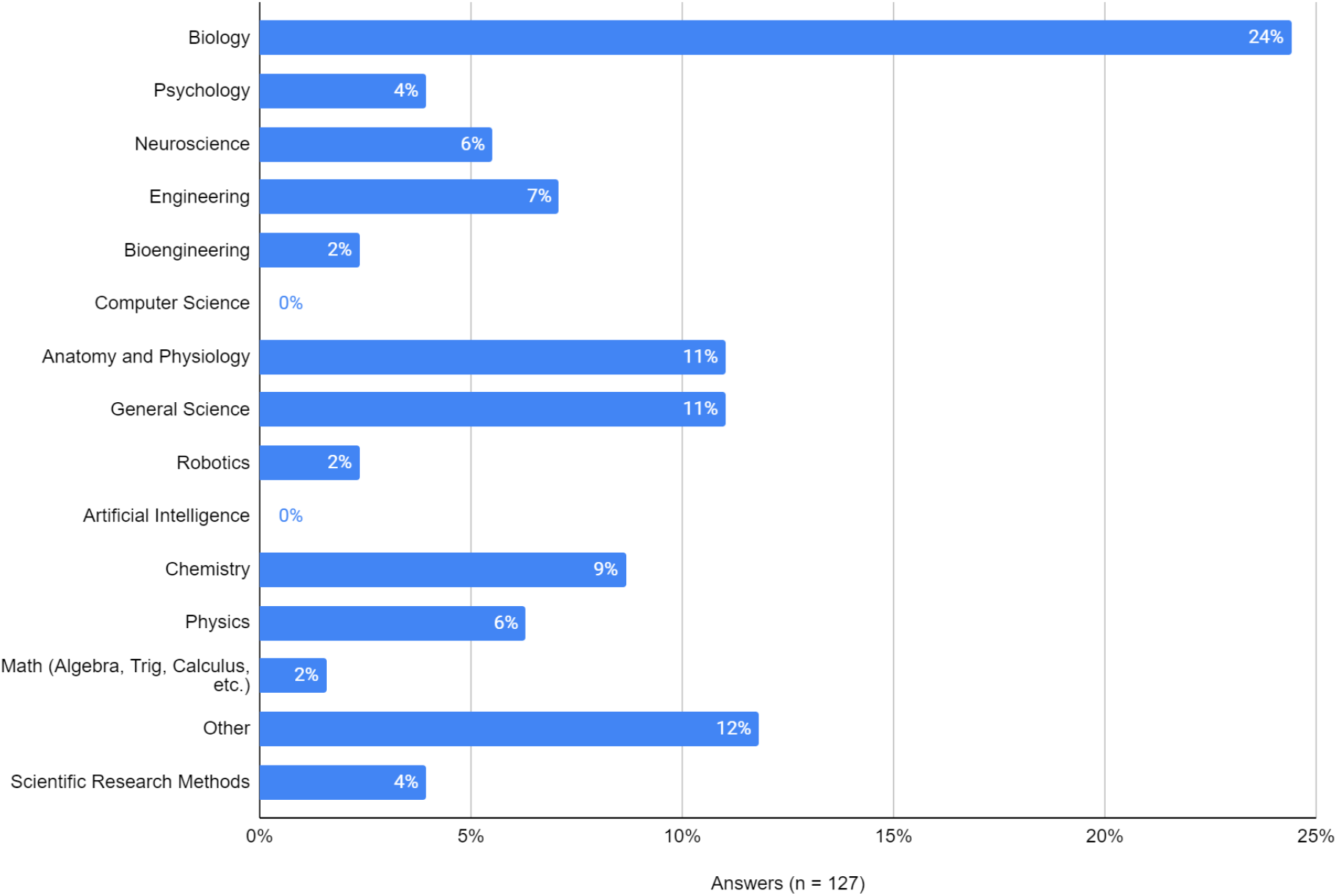

Q5. How much experience do you have doing scientific research? (select all that apply)

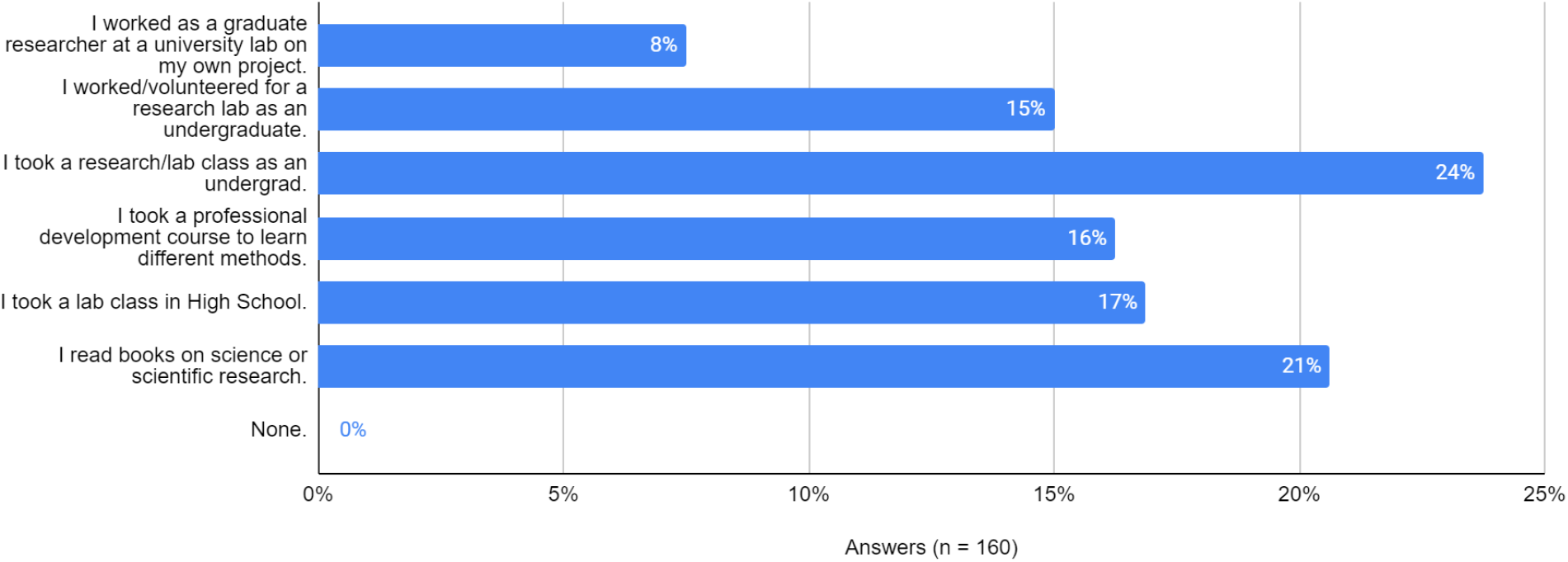

Q6. How comfortable do you feel with leading Phenomenon-based classroom activities where the student acts as a scientist and the outcomes of experiments are uncertain?

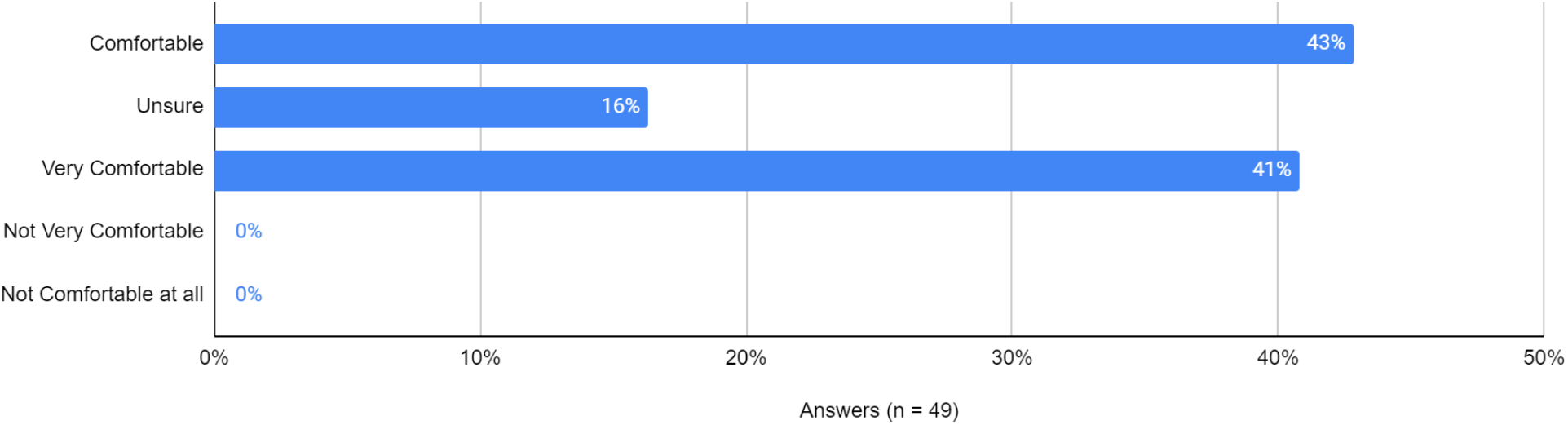

Q7. When working on group projects, how many groups do you typically have in your classroom?

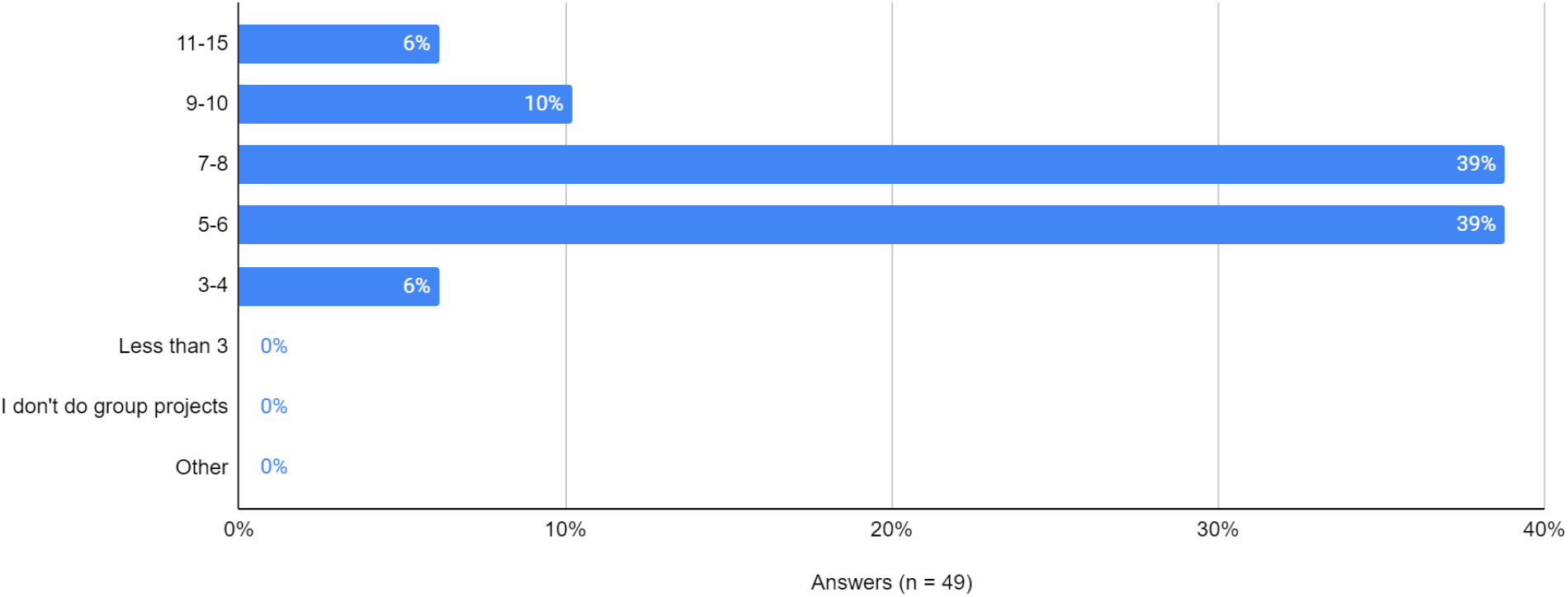

Q8. I find it easy to manage multiple groups working independently in the classroom?

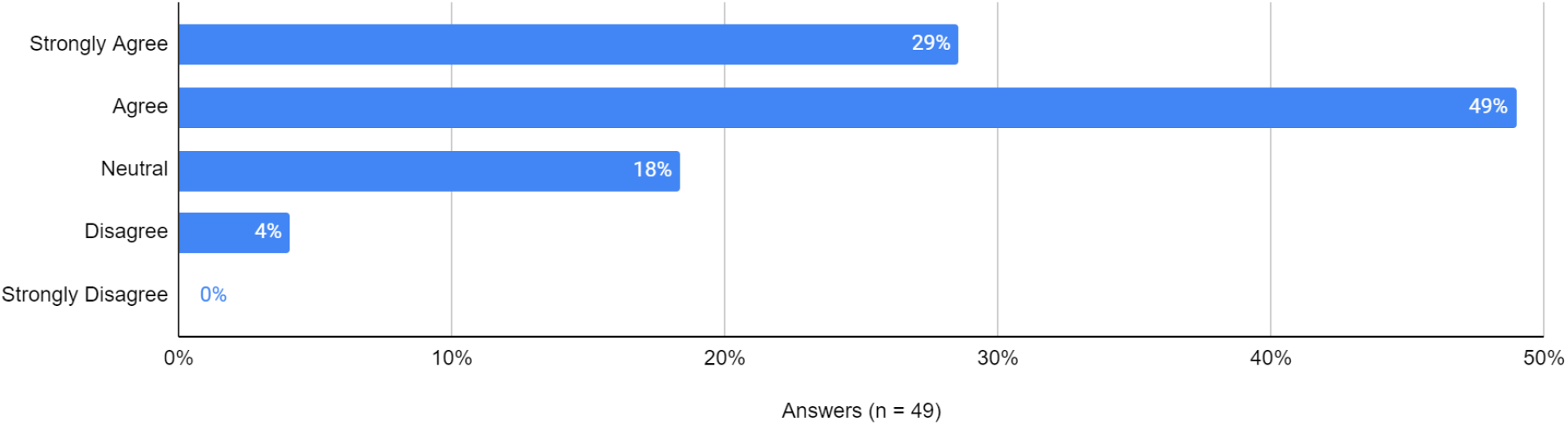

Q9. Would you be interested in PD to help develop your skills of leading independent scientific research skills in your classroom?

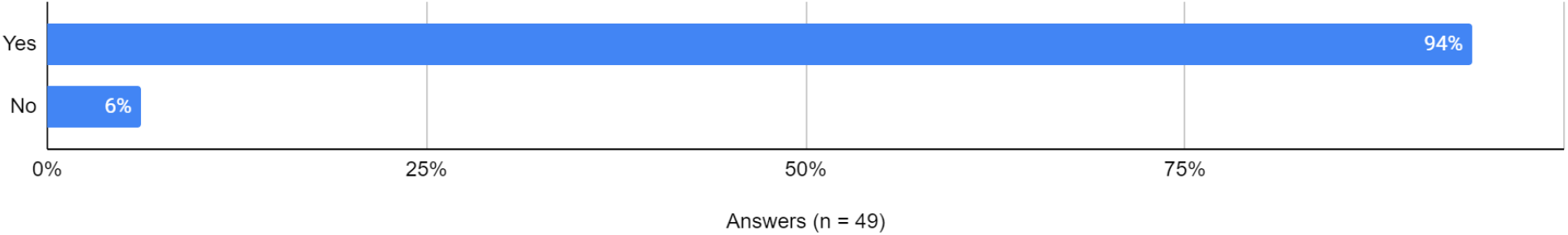

Q10. How many hours a month could you be available to participate in a paid, virtual (remote) professional development while you are still teaching classes in your home institution?

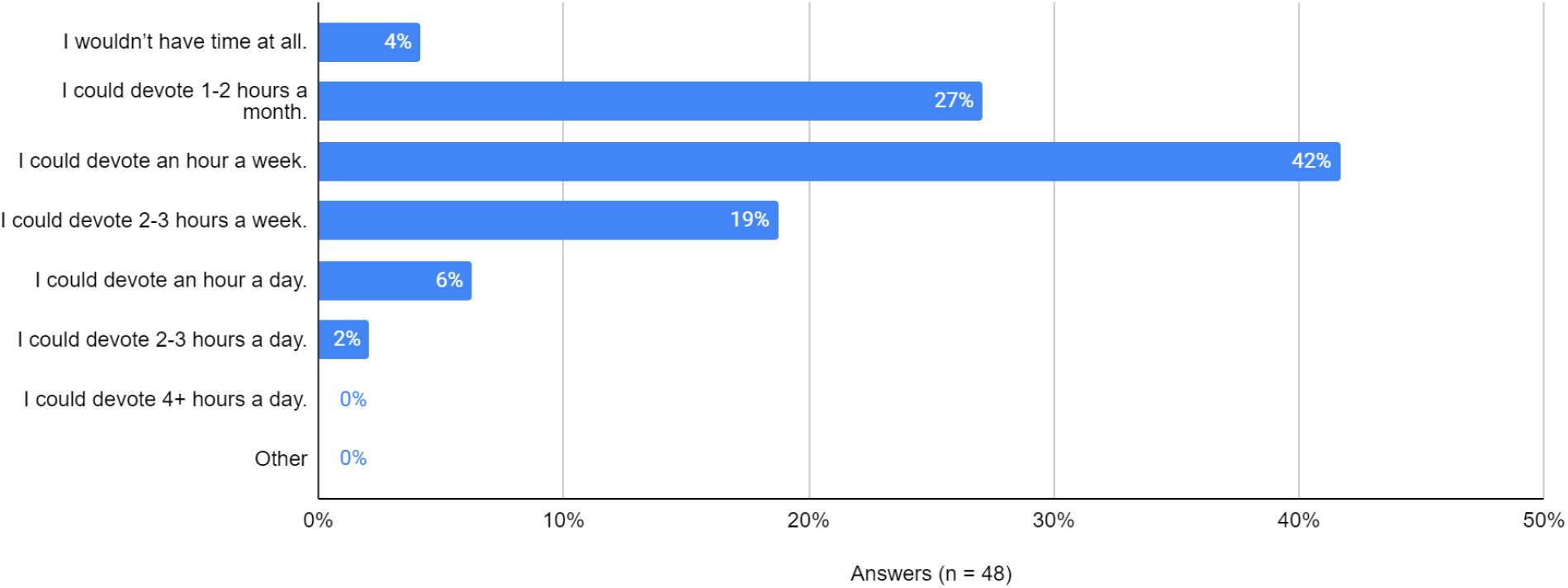

Q11. When does your school year end in the spring?

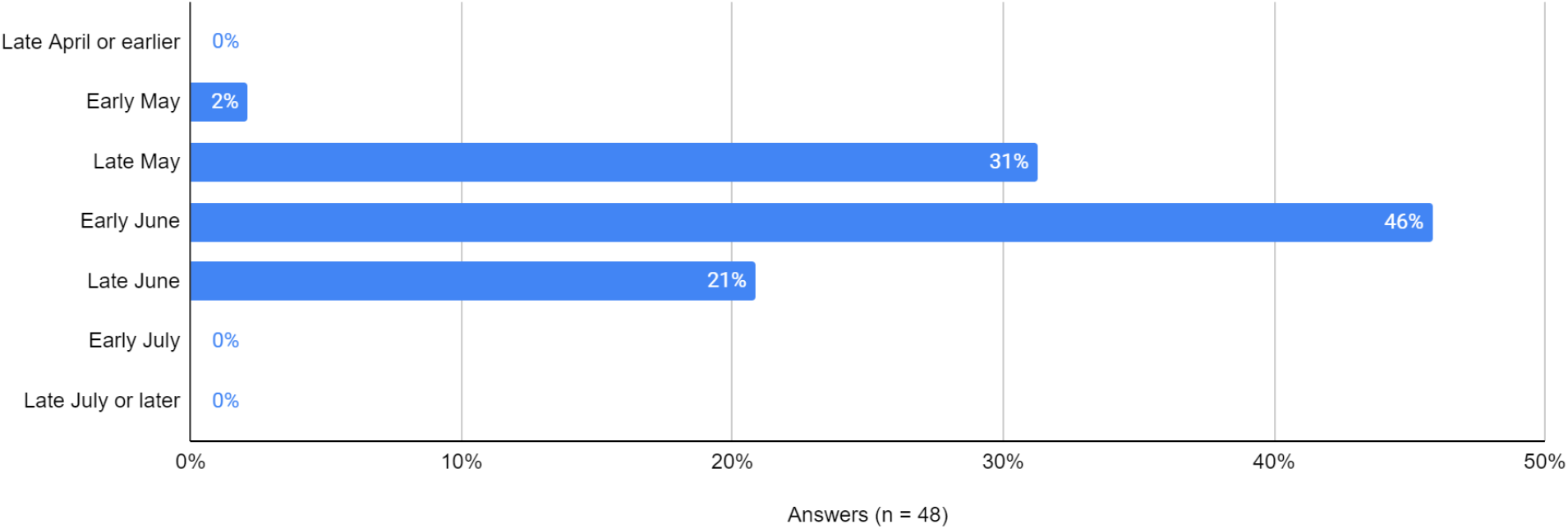

Q12. How many hours a month could you be available to participate in a paid, virtual (remote) PD when you are done teaching for the summer?

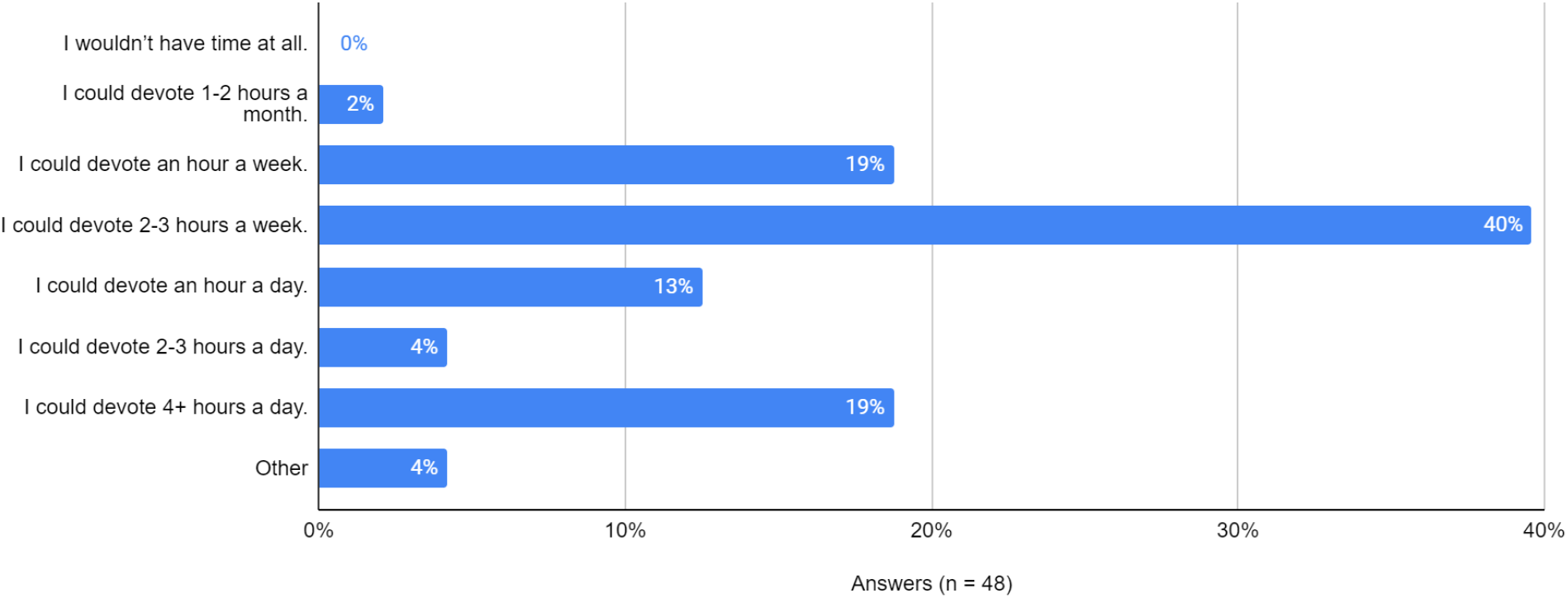

Q13. How interested would you be in participating in a virtual paid summer PD that lasts 10 weeks, a portion of which would start while you were still teaching in your home institution?

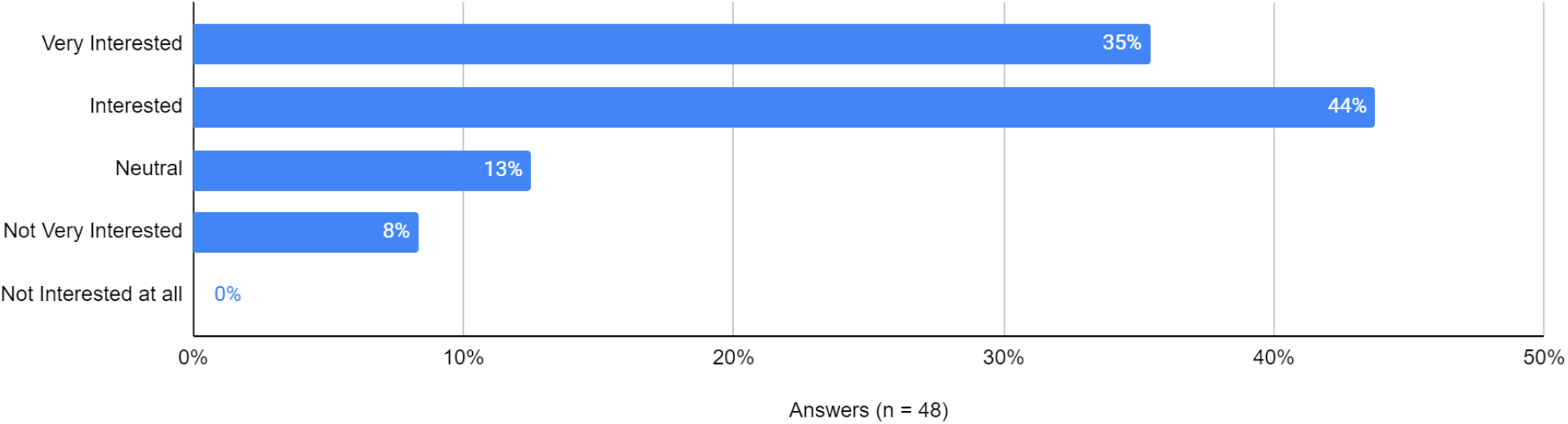

Q14. I would be interested in viewing an online weekly lecture series targeted at high school teachers that focused on (select all that apply).

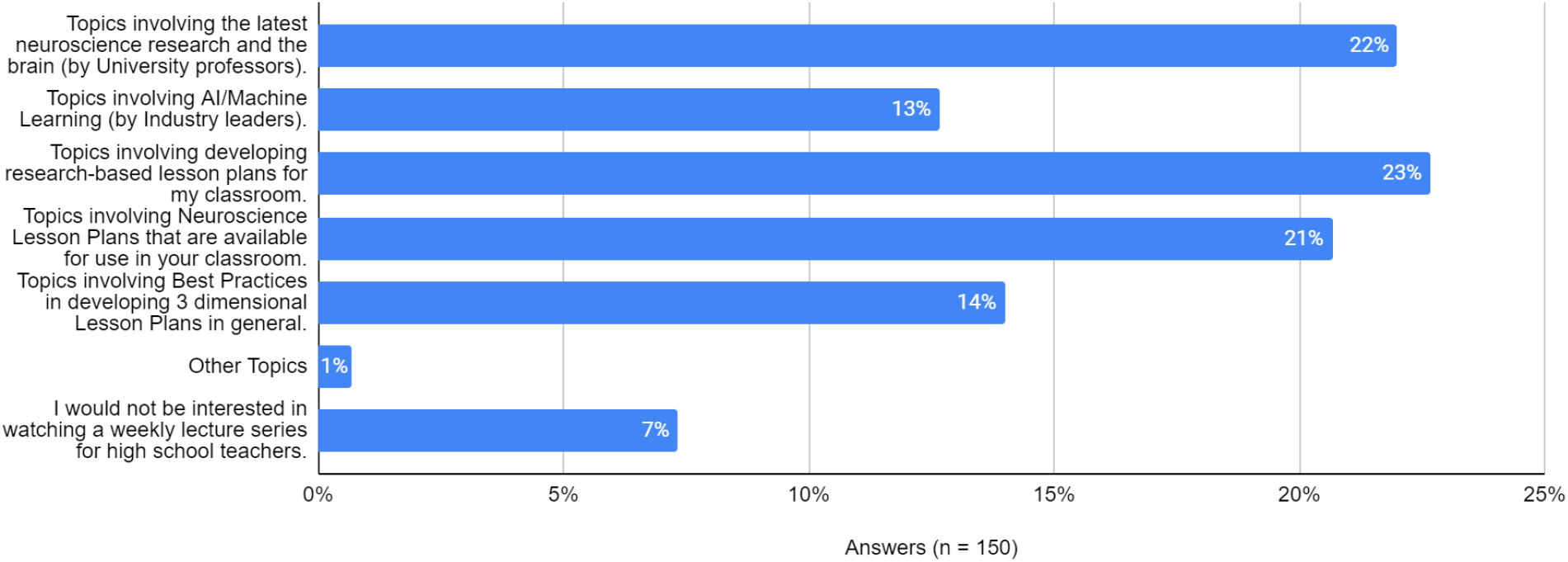

Q15. I develop Three Dimensional Learning (Scientific Practices, Core Ideas, and Cross-cutting concepts) lesson plans using the following method (select all that apply)

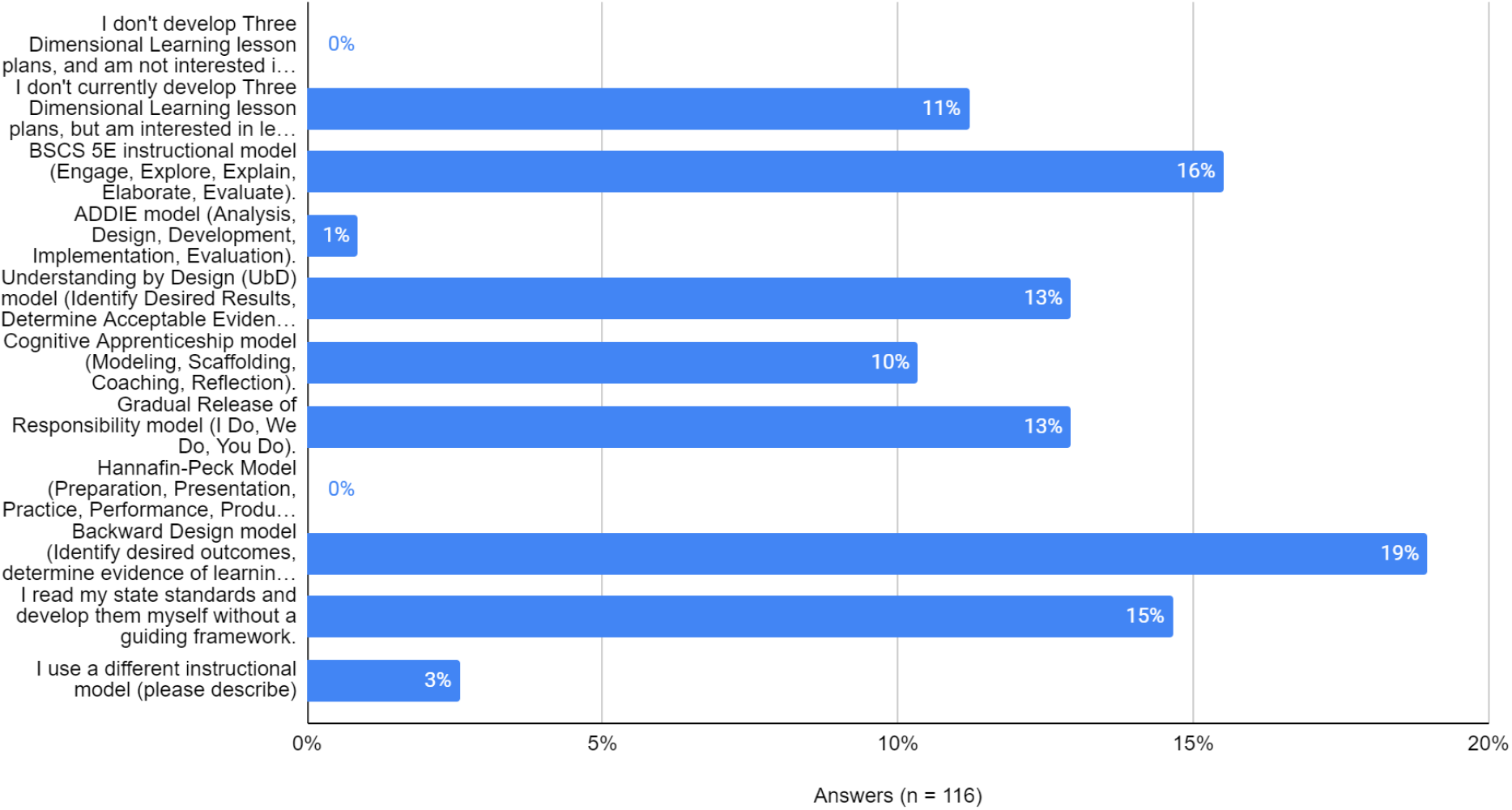

